# AS-Quant: Detection and Visualization of Alternative Splicing Events with RNA-seq Data

**DOI:** 10.1101/2020.02.15.950287

**Authors:** Naima Ahmed Fahmi, Hsin-Sung Yeh, Jae-Woong Chang, Heba Nassereddeen, Deliang Fan, Jeongsik Yong, Wei Zhang

## Abstract

A simplistic understanding of the central dogma falls short in correlating the number of genes in the genome to the number of proteins in the proteome. Post-transcriptional alternative splicing contributes to the complexity of proteome and are critical in understanding gene expression. mRNA-sequencing (RNA-seq) has been widely used to study the transcriptome and provides opportunity to detect alternative splicing events among different biological conditions. Despite the popularity of studying transcriptome variants with RNA-seq, few efficient and user-friendly bioinformatics tools have been developed for the genome-wide detection and visualization of alternative splicing events. We have developed AS-Quant (*A*lternative *S*plicing *Quant*itation), a robust program to identify alternative splicing events and visualize the short-read coverage with gene annotations. AS-Quant works in three steps: (i) calculate the read coverage of the potential splicing exons and the corresponding gene; (ii) categorize the splicing events into five different types based on annotation, and assess the significance of the events between two biological conditions; (iii) generate the short reads coverage plot with a complete gene annotation for user specified splicing events. To evaluate the performance, two significant alternative splicing events identified by AS-Quant between two biological contexts were validated by RT-PCR.

**Implementation:** AS-Quant is implemented in Python. Source code and a comprehensive user’s manual are freely available at https://github.com/CompbioLabUCF/AS-Quant

## 1 Introduction

A single gene can contain multiple exons and introns in eukaryotes. Exons can be joined together by splicing in different ways. Recent studies have estimated that alternative splicing events exist in more than 95% of multi-exon genes in human and mouse [10, 7, 1], and it provides cells with the opportunity to create protein isoforms with multiple functions from a single gene. Therefore, a precise detection of alternative splicing events among different biological contexts could provide insights into new molecular mechanisms and define high-resolution molecular signatures for phenotype predictions [8, 6]. High-throughput RNA-seq platform is capable of studying splicing variants, and several bioinformatics tools have been developed to identify alternative splicing events with RNA-seq [4, 2, 5, 9]. However, the selection for comprehensive and genome-wide assessments of the splicing events is limited, and few of the existing tools can provide high-resolution read coverage plots of the splicing events with accurate isoform annotation. We have developed AS-Quant, a program for genome-wide alternative splicing events detection and visualization. It efficiently handles large-scale alignment files with hundreds of millions of reads in different biological contexts and generates a comprehensive report for most, if not all, potential alternative splicing events, and generates high quality plots for the splicing events.

## 2 Methods

AS-Quant is composed of three steps: (i) read coverage estimation on exons; (ii) alternative splicing events categorization and assessment; (iii) visualization of splicing events (Figure 1). The first step requires aligned RNA-seq data in BAM format as input to estimate the read coverage on annotated exons and genes. In this step, AS-Quant generates read coverage files with SAMtools [3] to estimate the expression level for each exon and gene.

**Figure 1:**
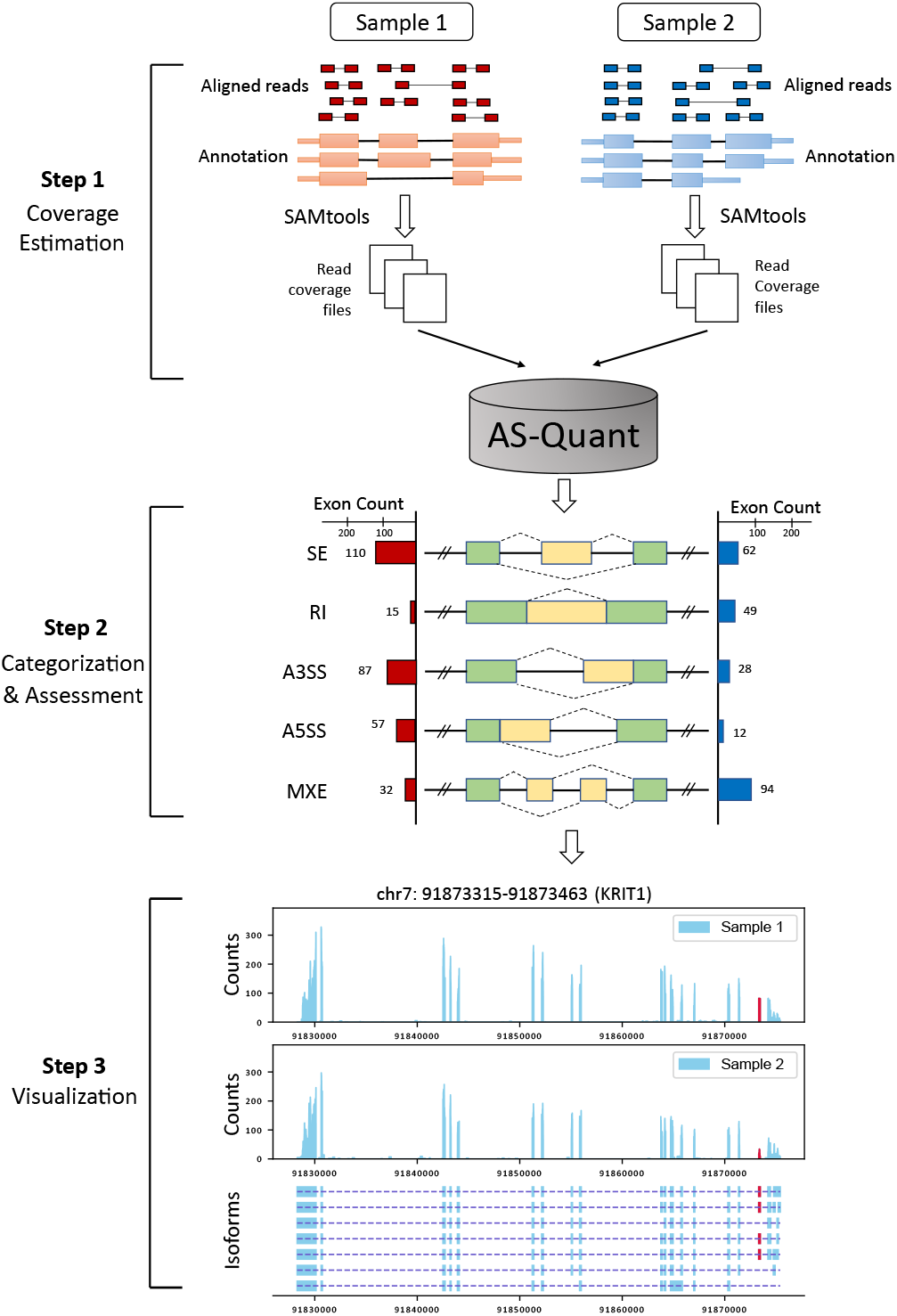
Workflow of AS-Quant. Starting with aligned RNA-seq bam files, AS-Quant consists of three steps (i) read coverage estimation, (ii) splicing events categorization and assessment, (iii) visualization.

In the second step, AS-Quant first identifies all potential alternative splicing events of five different categories based on RefSeq and UCSC gene annotation following the lead of the study in [4]. The five categories are: Skipped Exon (SE), Retained Intron (RI), Alternative 3’ Splice Site (A3SS), Alternative 5’ Splice Site (A5SS), and Mutually Exclusive Exon (MXE). The alternative splicing exon(s) in each category is highlighted in yellow in the middle panel of Figure 1. Then, AS-Quant calculates the average read coverage of the candidate alternative splicing exon (*n*) and all the other exons in the same gene (*N*) for both biological conditions. Next, the ratio differences between the two conditions are calculated based on 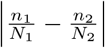, where 1 and 2 represent the two conditions. After that, a canonical 2 × 2 *χ*^2^-test is applied to report a *p*-value for each candidate splicing event. Only the alternative splicing events with a significant *p*-value (<0.1) and ratio difference (>0.1) will be reported in an Excel table. The horizontal bar charts in the middle panel of Figure 1 illustrate an example of number of the significant events in each category.

In the third step, based on the significant alternative splicing events reported in the second step, AS-Quant can generate RNA-seq read coverage plots with the isoform annotation for one or more user-specific events. An example is shown at the bottom panel in Figure 1. The alternative splicing exon is highlighted in red. In this step, users can enter the information of the chromosome region of the splicing exon from the output file in the second step and generate the RNA-seq read alignment plot.

## 3 Results

To evaluate the performance, AS-Quant was applied to RNA-seq data from two mouse embryonic fibroblasts (MEFs) samples, Tsc1−/− MEFs with control or U2af1 knocked down (KD). Based on the significant alternative splicing events between the two samples reported by AS-Quant, we generated the RNA-seq read coverage plots for two genes, *Ptbp1* and *Ganab*, as an example as shown in Figure 2(a). These genes were selected due to the design of PCR (polymerase chain reaction) primers for wet-lab validation. RT (reverse transcription)-PCR and agarose gel electrophoresis were applied to validate the expression of the transcript isoforms with exon inclusion/exclusion.

**Figure 2:**
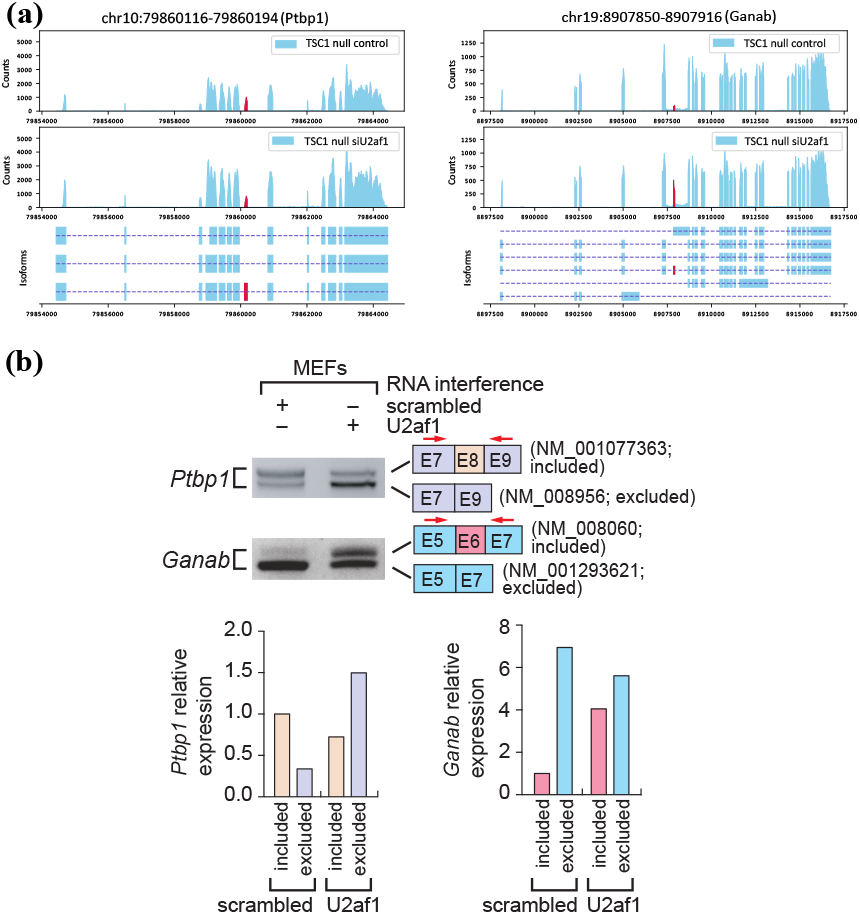
Experimental results: (a) RNA-seq read coverage plots of the gene *Ptbp1* and *Ganab* in the two samples with accurate isoform annotations. Alternatively spliced exons are marked in red. (b) Validation of isoform expression using RT-PCR and agarose gel electrophoresis. Quantitation of gel images using ImageQuant software is shown. Exon inclusion and exclusion events are color-coded. Total RNAs from mouse embryonic fibroblasts (MEFs) used for RNA-Seq experiments were used for RT-PCR amplification of *Ptbp1* or *Ganab* transcript isoforms. Scrambled RNA interference served as control and *U2af1* RNA interference is the case. The PCR primers to detect transcript isoforms for *Ptbp1* or *Ganab* are marked in red arrows. Alternative spliced isoform structures for each PCR product are shown. Exon numbers and transcript identification numbers in RefSeq annotation are shown. A higher band intensity of PCR products indicates a higher expression of that specific transcript isoform.

As shown in Figure 2(b), the relative expressions of the isoforms with exon inclusion/exclusion between the two samples showed significant changes, which is consistent with our observations on the RNA-seq read coverage plots reported in Figure 2(a). These results further confirm that AS-Quant can identify the true alternative splicing events in the RNA-seq samples from two different biological contexts.

The primer sequences used to measure the expression for transcript isoforms in the two genes are the following:

mPtbp1, forward 5’-TGCAGTATGCTGACCCTGTG-3’
mPtbp1 reverse 3’-AGCTGCACACTCTGATGCTT-5’
mGanab forward 5’-GATCGATGAGCTAGAGCCCC-3’
mGanab reverse 3’-TCCAAACCTACAGACGTGGG-5’

## 4 Conclusion

We present AS-Quant, a computational pipeline that allows the identification of transcriptome-wide alternative splicing events in RNA-seq data. The significant events are illustrated by read coverage plots along with full annotations of a specific gene. The experimental results on two mouse MEFs samples by RT-PCR demonstrate that AS-Quant is an accurate and efficient tool to detect alternative splicing events between samples with different biological contexts.

## User Manual

### About

AS-Quant is a computational tool used to detect alternative splicing(AS) events from RNA-seq data of two biological conditions (two samples). It can categorize five major types of AS in a comparative and comprehensive manner. AS-Quant also includes a visualization tool which generates plots for both the AS events and the annotation of the whole gene.

### Download

AS-Quant tool can be downloaded directly from github AS-Quant. Users need to have python installed on their machine. It can work on the Windows, Linux and Mac platform.

### Required softwares

1. Python (version 3.0 or higher)
2. Samtools 0.1.8 *[This specific version]

### Required python packages

1. matplotlib
2. scipy
3. pandas

### Run AS-Quant

AS-Quant is designed for handling both human and mouse Alternative Splicing events. The supplementary data (the five types of Alternative Splicing target dataset and the annotation) is provided in the project directory.

Users have to run the following two python files in order to run AS-Quant:

1. as quant.py: the main function which the users need to run
2. make plots.py: generates figures for visual representation of data

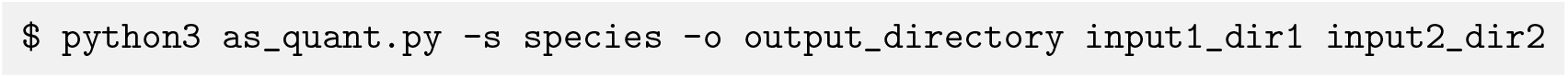

**Example**

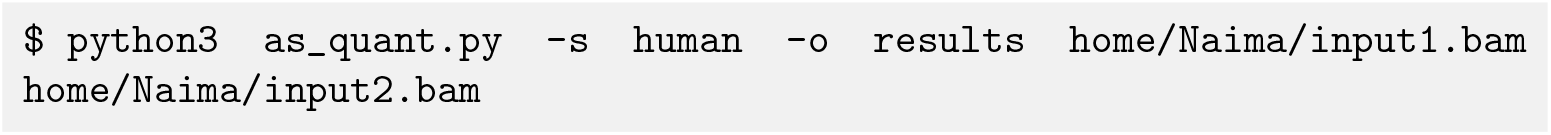

**Options:** (* refers to mandatory field)

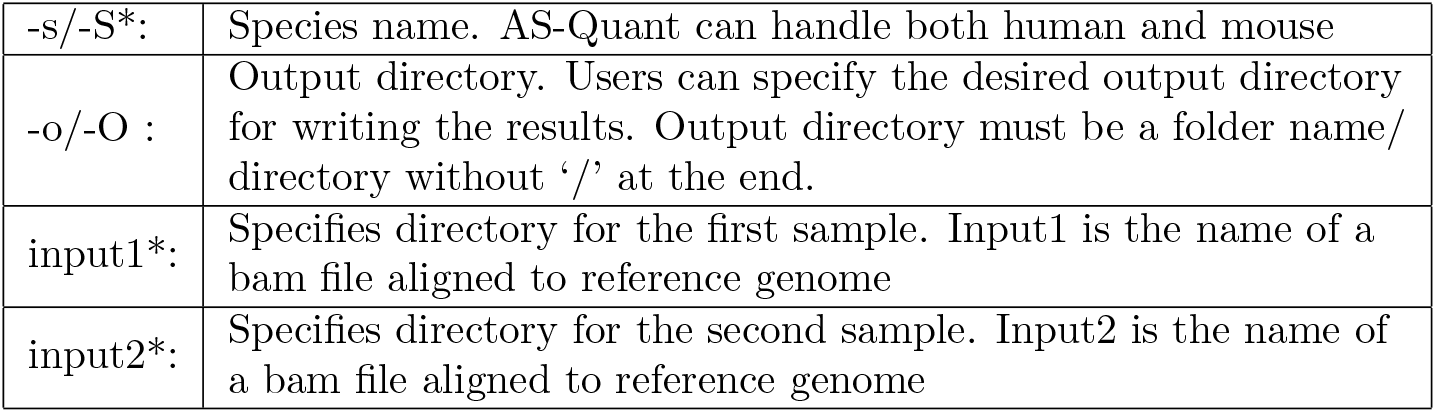

**as quant.py** will generate several intermediary files in the directory named **Output**(if the user does not provide a new output directory). After computing the significance of the association between the two samples, the final results will be written in the file named **sample1 Vs sample2.csv**. The following image is showing some of the generated fields in **sample1 Vs sample2.csv**:

|  | A | B | C | D | E | F | G | H |
| --- | --- | --- | --- | --- | --- | --- | --- | --- |
| 1 | Chrom | Gene Name | Exon Start | Exon End | p-value | Ratio difference | Absolute Ratio difference | Chrom region |
| 2 | chr1 | DPH5 | 101467022 | 101467100 | 0.9121302401 | 0.1000592552 | 0.1000592552 | chr1:DPH5:101467022-101467100 |
| 3 | chr1 | DPH5 | 101479265 | 101479374 | 0.5834973654 | 0.1790025326 | 0.1790025326 | chr1:DPH5:101479265-101479374 |
| 4 | chr1 | APITD1-CORT | 10494713 | 10494747 | 0.3359058847 | -0.972067033 | 0.972067033 | chr1:APITD1-CORT:10494713-10494747 |
| 5 | chr1 | PEX14 | 10596269 | 10596354 | 0.9023390528 | 0.137534426 | 0.137534426 | chr1:PEX14:10596269-10596354 |
| 6 | chr1 | PEX14 | 10659294 | 10659423 | 0.6931817062 | 0.2498379422 | 0.2498379422 | chr1:PEX14:10659294-10659423 |
| 7 | chr1 | AMPD2 | 110167924 | 110168055 | 0.563988963 | 0.1287839664 | 0.1287839664 | chr1:AMPD2:110167924-110168055 |
| 8 | chr1 | SLC16A4 | 110924273 | 110924417 | 0.5841909446 | -0.8237182045 | 0.8237182045 | chr1:SLC16A4:110924273-110924417 |
| 9 | chr1 | SLC16A4 | 110925455 | 110925588 | 0.6872561265 | -0.5067973124 | 0.5067973124 | chr1:SLC16A4:110925455-110925588 |
| 10 | chr1 | ST7L | 113098489 | 113098640 | 0.9287630199 | 0.196174765 | 0.196174765 | chr1:ST7L:113098489-113098640 |
| 11 | chr1 | ST7L | 113140592 | 113140708 | 0.7976173158 | 0.366711548 | 0.366711548 | chr1:ST7L:113140592-113140708 |
| 12 | chr1 | ST7L | 113143415 | 113143470 | 0.9416810826 | -0.1870431467 | 0.1870431467 | chr1:ST7L:113143415-113143470 |

### Run AS-Quant with provided sample input (Optional)

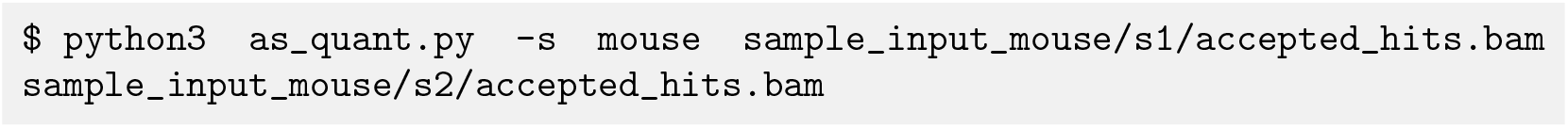

It will generate the output tables inside of folder ‘Output’ in the same directory. Or you can generate output in your desired directory, such as ‘Results’:

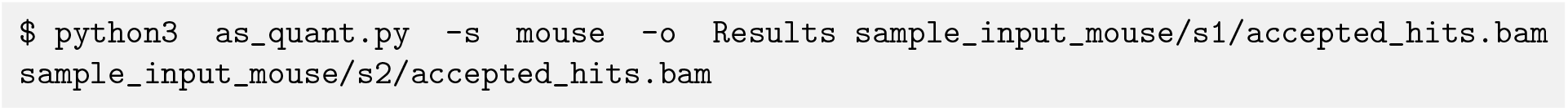

### Run make plots.py

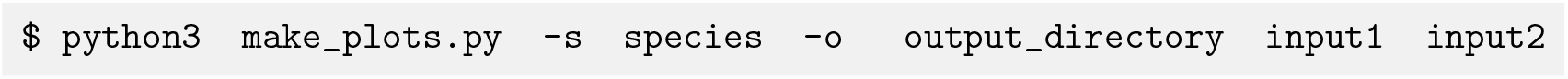

**Example:**

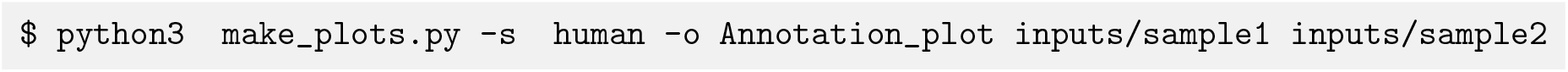

At that point, make plots.py will ask the user to enter the region of interest, for which they want to generate the annotation plot. The format should be in a specific format:

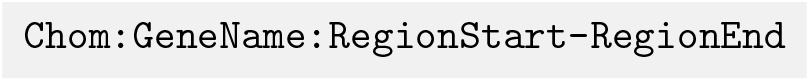

|  |  |
| --- | --- |
| Chrom : | Name of the chromosome |
| GeneName : | Name of the gene |
| RegionStart : | Starting position of the region |
| RegionEnd : | End position of the region |

**Example**

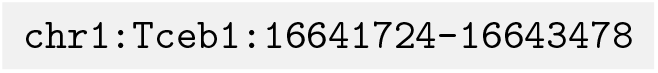

**make plots.py** will generate the read coverage plot for the given gene along with the whole annotation plot with all exons information of that gene. The output will produce a figure like the following:

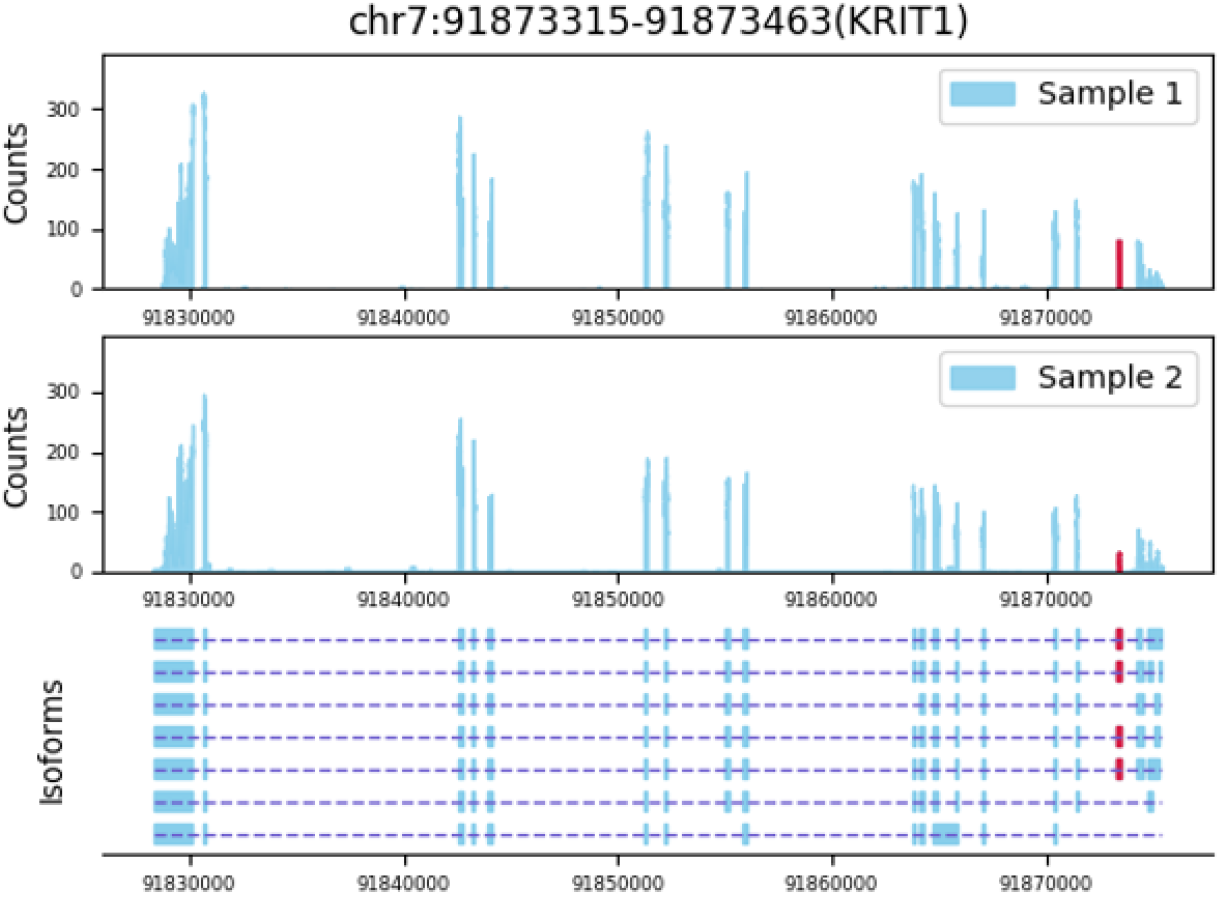

The first two subplots of the figure represent the read coverage of the two biological conditions. The bottom subplot shows the gene annotation along with all the exons information of that gene.

